# Conserved Brain-Associated CD8^+^ T Cell Programs and Their Regulation by the Brain Microenvironment

**DOI:** 10.64898/2026.10.22.761115

**Authors:** Kyungsoo Park, Tavian Sanchez, Shin Jeon, Peter F. Schade, David G. C. Hildebrand, Yeong Shin Yim

## Abstract

CD8^+^ T cells are increasingly recognized as components of the healthy brain, yet whether their specialized functional characteristics are conserved across species and how the brain microenvironment regulates their effector programs remain poorly understood. Here, we characterized brain-associated CD8^+^ T cells in the common marmoset and investigated their regulation by brain-derived signals. Flow cytometric analysis revealed a substantial population of CD8^+^ T cells in the healthy marmoset brain parenchyma that exhibited high IFN-γ competence but minimal granzyme B (GzmB) expression, resembling the functional phenotype previously identified in mice. Single-cell transcriptomic analyses of mouse and marmoset CD8^+^ T cells revealed tissue-associated transcriptional heterogeneity and distinct differentiation- and effector-associated programs. To investigate how the brain microenvironment shapes these characteristics, we exposed activated mouse CD8^+^ T cells to brain lysate and performed targeted CRISPR interference screening of 107 cell-surface receptor genes coupled with single-cell RNA sequencing. Brain lysate selectively reduced *Gzmb* and *Ifng* expression while increasing *Tnf* and *Xcl1*, indicating differential regulation of cytotoxic and inflammatory effector programs. Receptor perturbations revealed heterogeneous transcriptional responses and identified candidate pathways potentially involved in sensing brain-derived signals. Together, these findings identify shared functional characteristics of brain-associated CD8^+^ T cells across rodents and nonhuman primates and establish a framework for understanding how the neural microenvironment shapes adaptive immune function.

## Introduction

CD8^+^ T cells are classically defined by their ability to recognize peptide-MHC class I complexes and mount antigen-dependent responses, including cytokine production and cytotoxicity.^1–3^ However, their functional properties vary substantially across tissues.^4,5^ Within nonlymphoid organs, local cellular, molecular, and metabolic signals shape T cell differentiation and effector programs, allowing immune responses to adapt to the physiological requirements of individual tissues.^5–11^ Understanding this tissue-dependent regulation is particularly important in the central nervous system (CNS), where immune surveillance must coexist with the preservation of neural integrity.^12–15^ Whether CD8^+^ T cells acquire specialized functional characteristics within the healthy brain, and whether these features are conserved across species, remain important questions.

The brain was historically considered an immune-privileged organ,^16–19^ and studies of CNS-associated T cells have largely focused on infection, autoimmunity, injury, and neurodegeneration.^15,18,20–23^ Increasing evidence has expanded this view by identifying adaptive immune cells as components of the healthy brain and its border tissues.^24–30^ Our recent work identified a population of CD8^+^ T cells that becomes established in the healthy mouse brain during early postnatal development and progressively acquires an antigen-experienced phenotype.^31^ These cells contribute to neural homeostasis through an IFN-γ-dependent CD8^+^ T-cell–neuronal axis.^31^ Despite their differentiated phenotype and cytokine competence, brain-associated CD8^+^ T cells maintain low granzyme B (GzmB) expression, a major component of the canonical cytotoxic program.^31–33^ These observations suggest that cytokine competence and cytotoxic effector functions can be differentially regulated within the healthy brain,^24,31^ raising the possibility that the neural microenvironment supports a specialized form of adaptive immune regulation.

Whether these functional characteristics extend beyond rodents remains unknown. Differences in immune history, lifespan, and brain biology could influence both the composition and functional state of brain-associated T cells.^20,23–25,34–36^ Establishing which features are shared between mice and nonhuman primates is therefore essential for distinguishing broadly conserved tissue adaptations from species-specific immune characteristics. The common marmoset provides an opportunity to investigate this question by examining CD8^+^ T cells across brain parenchyma, brain-border tissues, and peripheral compartments.^37–39^ In particular, comparing the brain parenchyma with the dura may help distinguish functional characteristics associated with the neural environment from those shared with neighboring immune compartments.^25,28^

Beyond establishing the conservation of these characteristics, a central challenge is to understand how the brain microenvironment influences CD8^+^ T cell functional identity. Our previous finding that healthy brain lysate suppresses GzmB expression in activated peripheral CD8^+^ T cells indicates that brain-derived factors can directly modify an established effector program.^31^ However, whether these signals broadly suppress T-cell effector functions or selectively regulate individual functional programs remains unclear. Furthermore, the cell-surface receptors and signaling pathways through which CD8^+^ T cells respond to brain-derived factors remain poorly understood. Identifying these pathways may provide insight into how local tissue signals shape adaptive immune function in the healthy brain.

Here, we combine flow cytometry, single-cell transcriptomics, and targeted CRISPR interference (CRISPRi) screening to investigate the functional characteristics of brain-associated CD8^+^ T cells and their regulation by brain-derived signals. We first characterize CD8^+^ T cells in the healthy marmoset brain and compare their functional properties with those previously identified in mice. We then examine tissue-associated transcriptional heterogeneity and differentiation-related programs using single-cell RNA sequencing. Finally, we assess how brain lysate influences CD8^+^ T-cell effector programs and screen 107 candidate surface receptor genes to identify potential regulators of these responses. Together, these approaches investigate the conservation of brain-associated CD8^+^ T-cell functional characteristics across species and establish a framework for understanding how the neural microenvironment shapes adaptive immune cell identity.

## Results

### CD8^+^ T cells in the healthy marmoset brain exhibit cytokine competence with restrained granzyme B expression

Our recent work identified a distinct population of CD8^+^ T cells in the healthy mouse brain that exhibits antigen-experienced characteristics and high cytokine competence while maintaining low GzmB expression.^31^ To determine whether similar immune populations are present in nonhuman primates, we characterized brain-associated immune cells in common marmosets (*Callithrix jacchus*). To distinguish brain-associated immune cells from circulating leukocytes, animals received an intravenous injection of anti-marmoset CD45 antibody before perfusion. Following perfusion, the brain parenchyma and dura were collected separately, enzymatically dissociated, and subjected to Percoll density-gradient enrichment. Blood and spleen samples were collected for comparison, and immune populations were characterized by flow cytometry (Fig. 1A).

**Figure 1.**
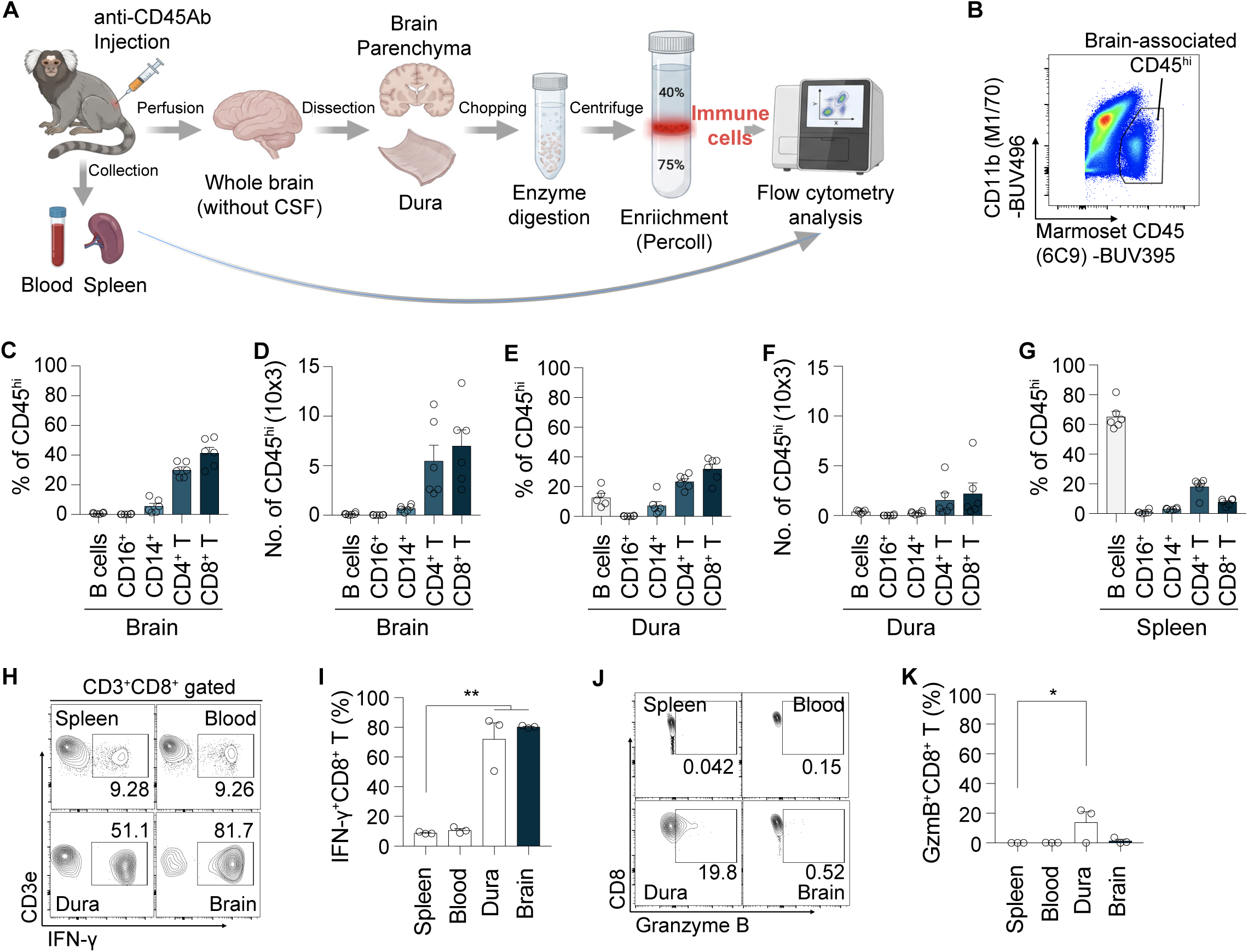
CD8^+^ T cells in the uninflamed marmoset brain demonstrate high cytokine competence alongside suppressed cytotoxic programming. **(A)** Experimental workflow for isolating and analyzing marmoset (*Callithrix jacchus*) immune subsets. To discriminate tissue-associated leukocytes from intravascular contamination, marmosets were intravenously injected with anti-CD45 antibody prior to cardiac perfusion. Brain parenchyma and dural tissue were subsequently harvested, enzymatically digested, and isolated via Percoll gradient centrifugation. Peripheral blood and splenic lymphocytes were processed in parallel. **(B)** Gating strategy and flow cytometric profile defining brain-resident CD45^hi^ immune populations according to CD45 and CD11b expression. (**C and D**) Proportional distribution (**C**) and total yield (**D**) of B cells, CD16^+^ subsets, CD14^+^ monocyte lineages, CD4^+^ T cells, and CD8^+^ T cells within the CD45^hi^ compartment of the brain parenchyma. (**E and F**) Lineage proportions (**E**) and cell counts (**F**) of specified immune compartments among dural CD45^hi^ leukocytes. (**G**) Relative abundance of designated immune subsets within splenic CD45^hi^ leukocytes. (**H and I**) Cytometry panels (**H**) and aggregate summary (**I**) showing IFNγ production by CD3^+^CD8^+^ T cells recovered from spleen, circulation, dural meninges, and parenchymal tissue. (**J and K**) Flow cytometric profiles (**J**) and cumulative data (**K**) indicating GzmB expression within CD3^+^CD8^+^ T cells harvested from the indicated tissue compartments. Data points denote individual animals. Histograms indicate mean ± SEM.

Flow cytometric analysis identified a distinct CD45^hi^ immune cell population in the marmoset brain (Fig. 1B). Within the parenchymal CD45^hi^ compartment, CD8^+^ T cells represented the largest immune cell population (∼40%), followed by CD4^+^ T cells (∼30%), whereas B cells, CD16^+^ cells, and CD14^+^ cells constituted smaller fractions (Fig. 1C). Absolute quantification further demonstrated that CD8^+^ and CD4^+^ T cells were major components of the parenchymal immune compartment (Fig. 1D). T cells were also enriched in the dura, although their relative proportions and absolute numbers differed from those in the brain parenchyma (Fig. 1E,F). In contrast, splenic CD45^hi^ immune cells were predominantly composed of B cells, with comparatively smaller proportions of CD8^+^ and CD4^+^ T cells (Fig. 1G). These findings demonstrate that the healthy marmoset brain contains a distinct immune cell composition characterized by a substantial representation of T lymphocytes.

We next examined the functional properties of CD8^+^ T cells across these compartments by assessing IFN-γ and GzmB expression. Intracellular cytokine staining revealed that CD8^+^ T cells isolated from the brain parenchyma and dura exhibited substantially higher IFN-γ competence than their circulating and splenic counterparts (Fig. 1H). Approximately 80% of parenchymal and 70% of dural CD8^+^ T cells were IFN-γ-positive, compared with approximately 10% of CD8^+^ T cells from blood and spleen (Fig. 1I). Despite this pronounced cytokine competence, GzmB expression remained minimal in brain parenchymal CD8^+^ T cells (Fig. 1J,K). Notably, dural CD8^+^ T cells exhibited a higher frequency of GzmB expression than parenchymal CD8^+^ T cells, despite similarly elevated IFN-γ competence. These observations reveal a functional distinction between CD8^+^ T cells in the brain parenchyma compared to those in the adjacent meningeal compartment.

Together, these findings demonstrate that CD8^+^ T cells in the healthy marmoset brain exhibit high IFN-γ competence coupled with restrained GzmB expression. This functional phenotype resembles that previously observed in the healthy mouse brain, suggesting that the differential regulation of cytokine competence and cytotoxic effector molecules represents a shared feature of brain-associated CD8^+^ T cells across mammalian species.

### Single-cell transcriptomics reveals tissue-associated CD8^+^ T cell states and distinct effector programs

To further characterize the molecular features of brain-associated CD8^+^ T cells, we performed single-cell RNA sequencing (scRNA-seq) of CD8^+^ T cells isolated from the marmoset brain parenchyma, dura, and peripheral compartments. Comparative analysis incorporating mouse CD8^+^ T-cell datasets^31^ enabled examination of transcriptional heterogeneity across anatomical compartments and species. Uniform Manifold Approximation and Projection (UMAP) visualization revealed heterogeneous CD8^+^ T-cell populations with tissue-associated differences in their transcriptional distributions (Fig. 2A). Brain parenchymal CD8^+^ T cells occupied transcriptional regions that differed from those of peripheral populations, suggesting that anatomical location is associated with distinct CD8^+^ T-cell states.

**Figure 2.**
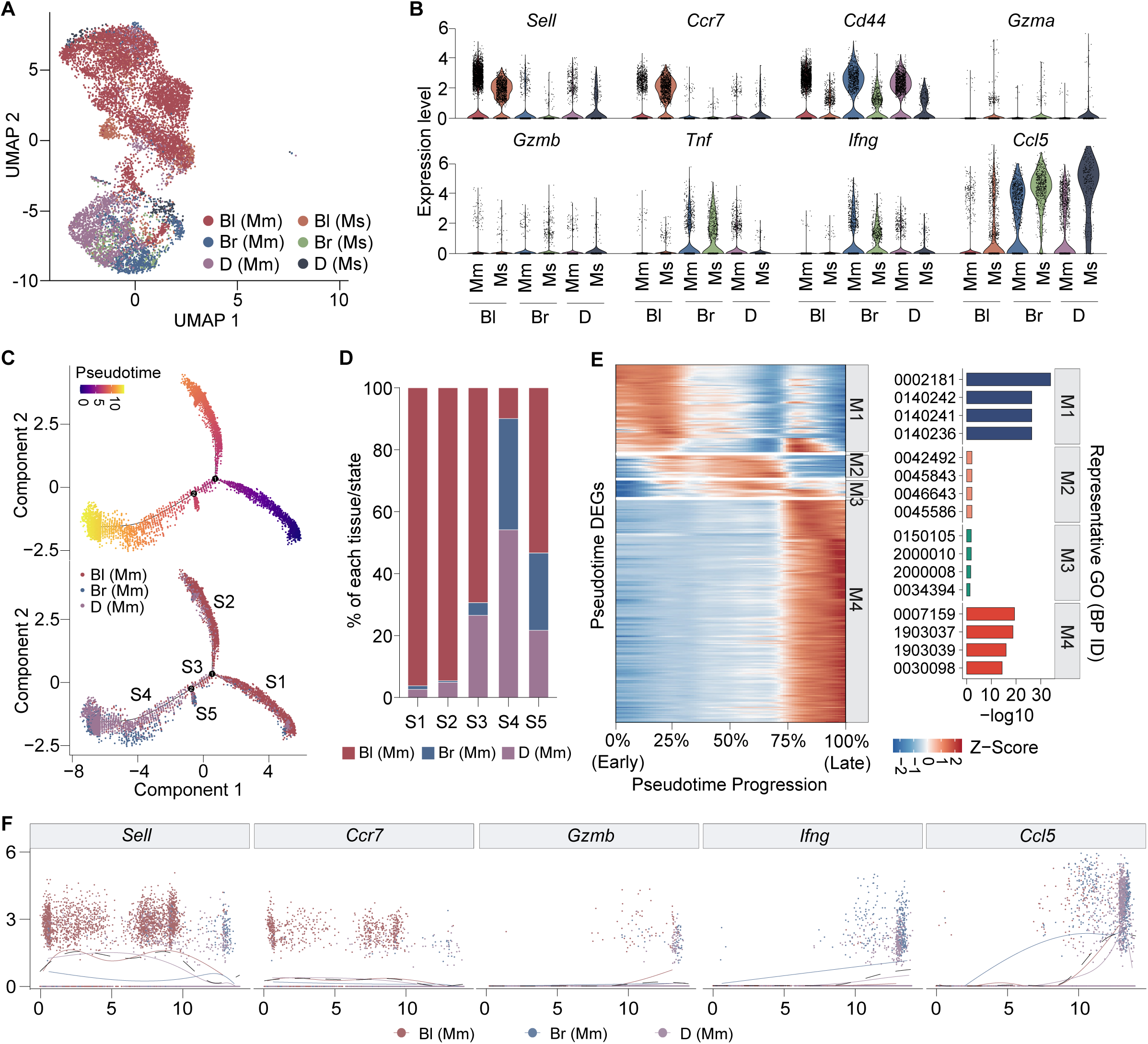
Single-cell transcriptomic analysis reveals tissue-associated transcriptional specialization of CD8^+^ T cells in the marmoset brain. **(A)** UMAP visualization of CD8^+^ T cells from marmoset (Mm) and mouse (Ms) blood (Bl), brain parenchyma (Br), and dura (D), colored by species and tissue origin. **(B)** Violin plots showing the expression of representative naïve-associated (*Sell*, *Ccr7*), activation-associated (*Cd44*), and effector-associated (*Gzma*, *Gzmb*, *Tnf*, *Ifng*, and *Ccl5*) genes in marmoset and mouse CD8^+^ T cells across the indicated tissue compartments. **(C)** Pseudotime trajectory analysis of marmoset CD8^+^ T cells isolated from blood, brain parenchyma, and dura. Upper panel shows cells colored by inferred pseudotime progression. Lower panel shows the distribution of cells from each tissue compartment along the inferred trajectory, with five transcriptional states (S1–S5) indicated. **(D)** Relative contributions of marmoset CD8^+^ T cells from blood, brain parenchyma, and dura to each of the five pseudotime states (S1–S5). **(E)** Heatmap showing genes with dynamic expression across the inferred pseudotime trajectory, organized into four transcriptional modules (M1–M4). Expression values are displayed as row-scaled Z-scores. Right, representative Gene Ontology (GO) enrichment results for each module, plotted as −log10. **(F)** Expression dynamics of selected naive-associated (*Sell*, *Ccr7*, *Tcf7*) and effector-associated (*Gzmb*, *Gzmk*, *Ifng*, *Ccl5*) genes along the inferred pseudotime trajectory. Points represent individual cells, colored by tissue origin. Curves indicate fitted expression trends.

We next examined the expression of genes associated with T-cell differentiation and effector functions. Naive-associated genes, including *Sell* and *Ccr7*, and effector-associated genes, including *Gzmb*, *Ifng*, and *Ccl5*, exhibited heterogeneous expression patterns across CD8^+^ T-cell populations (Fig. 2B). Consistent with our flow cytometric findings, brain-associated CD8^+^ T cells exhibited limited *Gzmb* expression, supporting the presence of a restrained GzmB-associated cytotoxic program. In contrast, other differentiation- and effector-associated transcripts displayed distinct expression patterns across compartments. These observations suggest that the specialized transcriptional characteristics of brain-associated CD8^+^ T cells extend beyond the regulation of individual cytotoxic molecules.

To further investigate the relationships among these transcriptional states, we performed pseudotime trajectory analysis. This analysis revealed a branched transcriptional continuum containing distinct cell populations distributed along the inferred trajectory (Fig. 2C). Examination of anatomical origins demonstrated that CD8^+^ T cells from different tissue compartments were not uniformly distributed across the trajectory, suggesting associations between tissue localization and transcriptional state (Fig. 2D). Analysis of gene expression along pseudotime identified four co-expression modules (M1–M4) with distinct expression patterns (Fig. 2E and Table 1). These modules suggest that multiple transcriptional programs contribute to the heterogeneity of CD8^+^ T-cell states. Examination of representative genes, including *Sell*, *Ccr7*, *Gzmb*, *Ifng*, and *Ccl5*, further revealed differential expression patterns along the inferred trajectory (Fig. 2F). Notably, *Gzmb* remained relatively limited across much of the trajectory, whereas other effector-associated genes exhibited more variable expression patterns. These findings suggest that individual components of CD8^+^ T-cell effector programs may be differentially regulated across transcriptional states.

Together, these analyses reveal substantial transcriptional heterogeneity among CD8^+^ T cells across brain-associated and peripheral compartments. The tissue-associated distributions and differential expression of effector-related genes support the presence of specialized CD8^+^ T-cell states within the brain, complementing the functional characteristics identified by flow cytometry. These findings provide a transcriptional framework for investigating how local tissue environments contribute to the functional specialization of brain-associated CD8^+^ T cells.

### Brain-derived signals selectively remodel CD8^+^ T cell effector programs through surface receptor-associated pathways

To investigate how CD8^+^ T cells sense and respond to signals within the brain microenvironment, we performed a targeted CRISPRi screen coupled with scRNA-seq. We generated a lentiviral dual-sgRNA library targeting 107 cell-surface receptor genes and transduced activated CD8^+^ T cells expressing dCas9 at a low multiplicity of infection. Transduced cells were subsequently cultured in the presence or absence of brain lysate, followed by scRNA-seq with sgRNA detection to assess receptor-dependent transcriptional responses (Fig. 3A).

**Figure 3.**
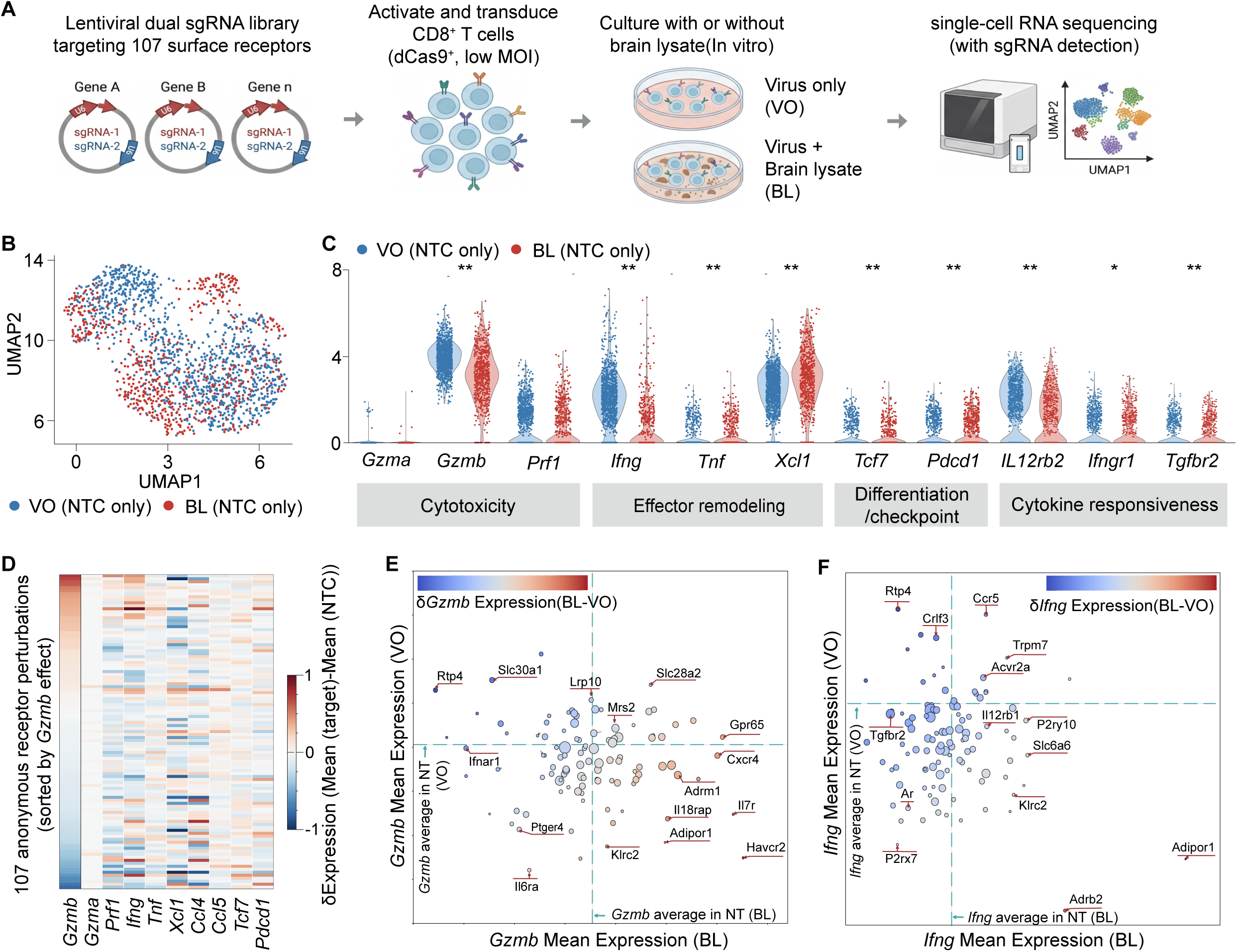
Brain-derived signals selectively remodel CD8^+^ T-cell effector programs through surface receptor-associated pathways. **(A)** Schematic of the targeted CRISPRi screening strategy. Activated dCas9-expressing CD8^+^ T cells were transduced at a low multiplicity of infection with a lentiviral dual-sgRNA library targeting 107 cell-surface receptor genes. Transduced cells were cultured in the absence (virus only, VO) or presence of brain lysate (BL), followed by scRNA-seq with sgRNA detection to assess receptor-dependent transcriptional responses. **(B)** Uniform manifold approximation and projection (UMAP) visualization of CD8^+^ T cells carrying non-targeting control (NTC) sgRNAs under VO (blue) and BL (red) conditions. **(C)** Violin plots showing the expression of representative genes associated with cytotoxicity (*Gzma, Gzmb, Prf1*), inflammatory effector functions (*Ifng, Tnf, Xcl1*), differentiation and checkpoint signaling (*Tcf7, Pdcd1*), and cytokine responsiveness (*Il12rb2, Ifngr1, Tgfbr2*) in NTC CD8^+^ T cells under VO (blue) and BL (red) conditions. Each dot represents an individual cell. Asterisks indicate statistically significant differences between conditions. Two-sided Mann–Whitney U tests * p<0.05, ** p<0.01 **(D)** Heatmap showing transcriptional responses across 107 cell-surface receptor perturbations under BL conditions. Rows represent individual receptor perturbations, and columns represent genes associated with cytotoxicity, inflammatory effector functions, and T-cell differentiation. Colors indicate relative changes in gene expression compared with NTC cells under BL conditions. **(E)** Screening-wide analysis of *Gzmb* expression across individual receptor perturbations under VO and BL conditions. Each dot represents one receptor perturbation, plotted according to mean *Gzmb* expression under BL (x-axis) and VO (y-axis) conditions. Dashed vertical and horizontal lines indicate the corresponding mean expression levels in NTC cells under BL and VO conditions, respectively. Dot colors represent the difference in mean *Gzmb* expression between BL and VO conditions (BL − VO). The four quadrants distinguish receptor perturbations associated with higher or lower *Gzmb* expression relative to condition-matched NTC controls. **(F)** Screening-wide analysis of *Ifng* expression across receptor perturbations under VO and BL conditions, presented as in (E). Each dot represents one receptor perturbation, with mean *Ifng* expression under BL (x-axis) and VO (y-axis). Dashed lines indicate condition-matched NTC mean expression levels, and dot colors represent the difference in mean *Ifng* expression between BL and VO conditions (BL − VO).

We first examined the effects of brain lysate on CD8^+^ T cells carrying non-targeting control (NTC) sgRNAs. UMAP visualization revealed broadly overlapping transcriptional populations between virus-only (VO) and brain lysate-treated (BL) conditions, although differences in their distributions were observed (Fig. 3B). Analysis of genes associated with CD8^+^ T-cell effector functions revealed selective transcriptional changes following brain lysate exposure (Fig. 3C). Notably, brain lysate reduced *Gzmb* expression without significantly altering *Gzma* or *Prf1*, suggesting that brain-derived signals selectively regulate components of the cytotoxic machinery rather than uniformly suppressing cytotoxic effector genes. This selective regulation extended to inflammatory mediators, with reduced *Ifng* expression but increased *Tnf* and *Xcl1* expression. In addition, brain lysate increased the expression of *Tcf7* and *Pdcd1*, genes associated with T-cell differentiation and activation states, respectively. Several cytokine receptor genes were also differentially expressed, including reduced *Il12rb2* and increased *Ifngr1* and *Tgfbr2*. Together, these findings indicate that brain-derived signals reshape multiple aspects of CD8^+^ T-cell functional identity rather than inducing generalized suppression of effector programs.

We next investigated whether cell-surface receptors contribute to these brain lysate-induced transcriptional changes. Screening-wide analysis of the 107 receptor perturbations revealed heterogeneous transcriptional effects across genes associated with cytotoxicity, inflammatory responses, and T-cell differentiation (Fig. 3D). To further identify receptor pathways potentially involved in regulating CD8^+^ T-cell effector functions, we focused on *Gzmb* and *Ifng*, two effector molecules whose expression was reduced following brain lysate exposure. For each receptor perturbation, we compared mean *Gzmb* or *Ifng* expression under VO and BL conditions using the corresponding NTC populations as reference points (Fig. 3E,F). This analysis identified subsets of receptor perturbations associated with increased *Gzmb* expression relative to NTC cells under BL conditions, suggesting that these receptors may contribute to the regulation of GzmB expression in response to brain-derived signals (Fig. 3E). Similarly, analysis of *Ifng* revealed receptor perturbations associated with increased or decreased expression under BL conditions relative to NTC controls (Fig. 3F). Notably, some receptor perturbations exhibited opposing expression patterns between VO and BL conditions, suggesting that their regulatory effects may depend on the surrounding microenvironment. Together, these findings identify candidate surface receptor pathways involved in regulating cytotoxic and inflammatory effector programs in response to brain-derived signals.

Collectively, these results demonstrate that brain-derived signals selectively remodel CD8^+^ T-cell effector programs and identify candidate surface receptor pathways that may contribute to this regulation. The heterogeneous effects observed across receptor perturbations suggest that multiple receptor-associated mechanisms participate in shaping CD8^+^ T-cell functional responses to the brain microenvironment.

## Discussion

The healthy brain presents a unique environment for adaptive immune cells, where immune surveillance must be balanced against the preservation of neural integrity.^12,13,15^ Although CD8^+^ T cells are increasingly recognized as components of the healthy central nervous system,^24,25,31,40^ whether their specialized functional characteristics are conserved across species and how the brain microenvironment shapes these properties remain poorly understood. Here, we demonstrate that CD8^+^ T cells in the healthy marmoset brain exhibit a distinctive functional phenotype characterized by high IFN-γ competence and limited GzmB expression, resembling the phenotype previously identified in the mouse brain.^31^ Single-cell transcriptomic analyses further reveal tissue-associated CD8^+^ T-cell states and distinct differentiation-associated programs across brain and peripheral compartments. Finally, using an in vitro brain lysate system coupled with targeted CRISPRi screening, we show that brain-derived signals selectively remodel CD8^+^ T-cell effector programs and identify candidate surface receptor pathways potentially involved in this regulation. Together, these findings support the concept that the healthy brain provides a specialized environment that shapes adaptive immune cell identity and function.

A central finding of this study is the presence of CD8^+^ T cells with shared functional characteristics in the healthy brains of mice and nonhuman primates. Our previous work demonstrated that CD8^+^ T cells progressively accumulate in the mouse brain during postnatal development, acquire antigen-experienced and tissue-associated characteristics, and contribute to neural homeostasis.^31^ Despite their differentiated phenotype and capacity to produce inflammatory cytokines, these cells maintain low GzmB expression. Here, we extend these observations to the common marmoset, demonstrating that brain parenchymal CD8^+^ T cells similarly exhibit high IFN-γ competence with minimal GzmB expression. Importantly, this phenotype differs from that observed in peripheral compartments and, particularly for GzmB, from the dura, suggesting that the brain parenchymal environment may impose distinct constraints on CD8^+^ T-cell effector functions. The identification of similar characteristics in rodents and nonhuman primates suggests that this functional organization may represent a broadly conserved feature of brain-associated adaptive immunity rather than a mouse-specific phenomenon.

The coexistence of cytokine competence and restrained GzmB expression raises an important question regarding how CD8^+^ T-cell effector functions are coordinated within the healthy brain. Canonical CD8^+^ T-cell differentiation is frequently associated with the acquisition of inflammatory and cytotoxic effector programs^3^. However, our findings suggest that these functions can be regulated independently within the brain microenvironment. IFN-γ may contribute to communication between CD8^+^ T cells and surrounding neural or glial populations,^29^ consistent with our previous demonstration of an IFN-γ-dependent CD8^+^ T-cell–neuronal axis involved in maintaining neural homeostasis.^31^ In contrast, limiting GzmB expression may reduce the potential for cytotoxic injury in a tissue containing vulnerable and largely nonrenewable neuronal populations.^13,14^ Although low GzmB expression alone does not establish an absence of cytotoxic activity,^33,41^ the combination of cytokine competence and restrained GzmB suggests that brain-associated CD8^+^ T cells adopt a functional state compatible with immune surveillance and tissue preservation. Whether this balance is actively maintained throughout life or altered under inflammatory and pathological conditions remains an important question.

Our single-cell transcriptomic analyses further support the existence of specialized CD8^+^ T-cell states associated with the brain microenvironment. CD8^+^ T cells from brain parenchyma, dura, and peripheral compartments displayed distinct transcriptional distributions and differentiation-associated gene expression patterns. Pseudotime analyses identified multiple transcriptional states and coordinated gene expression modules, suggesting heterogeneity in the differentiation programs represented across anatomical compartments. Notably, the differential expression of naïve-associated markers, inflammatory mediators, and cytotoxic genes indicates that tissue-associated adaptation involves broader transcriptional changes rather than isolated suppression of GzmB. The distinct regulation of individual granzymes is particularly interesting, as it suggests that cytotoxic effector molecules may be controlled through partially independent mechanisms. Nevertheless, the inferred trajectories represent relationships among transcriptional states rather than direct evidence of developmental transitions.^42^ Future studies using longitudinal sampling or lineage-tracing approaches will be necessary to determine how these states arise and whether brain-associated CD8^+^ T cells undergo progressive differentiation after entering the parenchyma.

An important unresolved question is whether the distinctive characteristics of brain-associated CD8^+^ T cells reflect selective recruitment of particular T-cell populations, local adaptation following tissue entry, or a combination of both processes. Our previous mouse studies support the possibility that CD8^+^ T cells undergo phenotypic changes after entering the developing brain.^31^ The present findings extend this framework by demonstrating that brain-derived factors can directly modify the transcriptional programs of activated peripheral CD8^+^ T cells in vitro. Exposure to brain lysate selectively reduced *Gzmb* and *Ifng* expression while increasing *Tnf* and *Xcl1*, without uniformly suppressing other cytotoxic mediators such as *Gzma* and *Prf1*. Brain lysate also altered genes associated with differentiation, checkpoint signaling, and cytokine responsiveness. These findings indicate that brain-derived signals do not simply inhibit CD8^+^ T-cell activation but instead induce a selective reorganization of effector-associated transcriptional programs. Such regulation may allow CD8^+^ T cells to retain particular immune functions while limiting others that could be detrimental within the neural environment.

Interestingly, the effects of brain lysate on activated peripheral CD8^+^ T cells do not fully reproduce the functional characteristics observed in brain-associated CD8^+^ T cells in vivo. In particular, although brain-associated CD8^+^ T cells exhibit high IFN-γ competence, brain lysate reduced IFN-γ expression in vitro. This apparent difference suggests that the establishment of brain-associated CD8^+^ T-cell identity is more complex than the acute response to soluble brain-derived factors alone. The functional state of CD8^+^ T cells within the intact brain may reflect the combined influence of local cytokines, antigen recognition, cell–cell interactions, metabolic signals, and the duration of tissue residence. Moreover, the effects of these signals may depend on the differentiation and activation state of the responding T cells. Thus, our in vitro findings should be interpreted as evidence that brain-derived factors can regulate selected components of CD8^+^ T-cell function, rather than as a complete recapitulation of the brain-associated phenotype.

To begin identifying how CD8^+^ T cells detect brain-derived signals, we performed a targeted CRISPRi screen of 107 cell-surface receptor genes coupled with single-cell transcriptomic profiling. This approach allowed us to examine the effects of receptor perturbations on multiple effector-associated genes under control and brain lysate conditions. Screening-wide analyses revealed heterogeneous transcriptional responses, with receptor perturbations associated with distinct patterns of *Gzmb* and *Ifng* expression. Comparisons with condition-matched non-targeting controls identified candidate receptor pathways potentially involved in regulating these effector programs. These observations support the possibility that CD8^+^ T-cell responses to the brain microenvironment are influenced by multiple receptor-associated mechanisms rather than a single generalized inhibitory pathway. Importantly, the distinct responses observed across cytotoxic and inflammatory genes suggest that different components of the effector program may be regulated independently. Further studies will be required to establish which receptors directly mediate brain-derived signals, identify their relevant ligands, and determine whether these pathways operate within brain-associated CD8^+^ T cells in vivo.

Several limitations should be considered. First, although the mouse and marmoset comparisons reveal shared functional characteristics, cross-species conservation of specific molecular mechanisms remains to be established. Differences in immune history, developmental timing, and environmental exposure may influence the composition and differentiation states of brain-associated CD8^+^ T cells.^23,43^ Second, the in vitro brain lysate system contains a complex mixture of neural and non-neural components and does not preserve the spatial organization or cellular interactions of the intact brain. Consequently, the identities, concentrations, and physiological availability of the signals responsible for the observed transcriptional changes remain unknown. Third, the CRISPRi screen provides a framework for identifying candidate receptor-dependent regulatory pathways, but perturbation-associated transcriptional changes require independent validation and functional characterization. Finally, although the shared phenotype is consistent with a specialized adaptation to the healthy brain, the present CRISPRi screening study does not directly establish the physiological consequences of these CD8^+^ T-cell programs in nonhuman primates.

In summary, our findings identify shared functional characteristics of brain-associated CD8^+^ T cells across mice and nonhuman primates and demonstrate that brain-derived signals can selectively reshape CD8^+^ T-cell effector programs. The coexistence of cytokine competence and restrained GzmB expression suggests that the healthy brain supports a specialized form of adaptive immune regulation in which inflammatory and cytotoxic functions are differentially controlled. By integrating cross-species immune profiling with single-cell transcriptomics and targeted genetic screening, this study establishes a foundation for investigating how the neural microenvironment regulates adaptive immune cell identity.

Understanding these mechanisms may provide insight into how immune surveillance is maintained without compromising neural integrity and how disruption of this balance could contribute to neurological disease.

## Methods

### Animals

#### Marmosets

All experimental procedures involving marmosets were approved by The Rockefeller University Institutional Animal Care and Use Committee and were performed in accordance with guidelines from the United States National Institute of Health. Six marmosets were used in this study, four males aged 3.6–7.6 years and two females aged 7.0–7.7 years. Marmosets were sedated with alfaxalone (7-12 mg/kg IM), administered midazolam (0.5–1.0 mg/kg IM) and buprenorphine (0.005mg/kg IM), and maintained on isoflurane (0.5–2.0% IH) anesthesia. Following sedation, an IV catheter was placed. To label immune cells residing in the blood or vasculature, 45μg of anti-CD45 labeling antibody in saline was then injected through the catheter. After 10 min, ≤2.0mL blood was collected through the catheter. The marmosets were then trans-cardially perfused, with a euthanasia solution (Euthasol, >120 mg/kg IV) administered immediately after starting the perfusion. The brain, dura, and spleen were then dissected for subsequent experiments.

#### Mice

All experimental procedures involving mice were approved by and conducted in accordance with the guidelines of the Institutional Animal Care and Use Committee of the University of Pennsylvania. All mice used for in vitro primary CD8^+^ T-cell culture, lentiviral transduction, and brain lysate treatment were homozygous for the dCas9 allele (Jax #030000). All mice were 5.5-month-old males. Mice were housed in filter-topped cages at 22.2 °C and 52.1% humidity under a 12-hour light–dark cycle, with autoclaved food and water available *ad libitum*.

### Marmoset immune cell preparation

Blood was collected via the IV catheter as described above and stored at room temperature in EDTA tubes. Following sedation and anti-CD45 antibody labeling (anti-marmoset CD45 antibody, clone 6C9, BioLegend Cat. #250204). Animals were euthanized and perfused as described above, and brain, spleen, and dura mater were collected. The brain was carefully removed from the skull, and the dura mater was dissected from the inner surface of the skull. All dissected tissues were stored in RPMI 1640 medium containing 10% FBS with 25mM HEPES. <u>Brain tissue</u> was minced with scissors and transferred to glass vials containing digested buffer of RPMI 1640 containing 10% FBS, 1 mg/ml collagenase IV from Clostridium histolyticum (Sigma), and 50 µg/ml DNase I for 25 minutes at 37 °C with magnetic stirring. Samples were passed through a 70-µm cell strainer and washed. Cells were resuspended in 40% Percoll and layered over 75% Percoll, followed by centrifugation at 2,500 rpm for 20 minutes at room temperature without braking. The upper myelin layer was removed, and immune cells at the interface were collected and washed twice for further experiments. <u>Dura mater</u> was minced with scissors transferred into glass vials containing digestion buffer of RPMI 1640 supplemented with 10% FBS, 0.75 mg/ml collagenase IV, and 50 µg/ml DNase I. Samples were digested for 8 minutes at 37 °C with magnetic stirring and washed twice before further processing.<u>Spleen tissue</u> was mechanically dissociated between glass slides. Red blood cells were lysed with RBC Lysis Buffer (Cell Signaling Technology, Cat. #46232) for 2 minutes at room temperature, and samples were passed through a 40-µm cell strainer and washed twice for further experiments. <u>Blood</u>, collected as described above, was processed by density-gradient centrifugation using Lymphoprep in SepMate-15 tubes (STEMCELL Technologies) to isolate peripheral blood mononuclear cells (PBMCs), following the manufacturer’s protocol. The PBMC-containing upper fraction was collected and washed twice for further experiments.

Single-cell suspensions were stained with the LIVE/DEAD Fixable Near-IR Dead Cell Stain Kit (Invitrogen, Cat. #L34993). Surface antigens were stained for 30 minutes at 4 °C using the following antibodies: anti-CD45 (Clone 6C9, BioLegend, Cat. No. 250206); anti-CD3 (Clone SP34-2, BD Horizon, Cat. No. 560352); anti-CD4 (Clone L200, BD Horizon, Cat. No. 563737); anti-CD8 (Clone HIT8a, BioLegend, Cat. No. 300910); anti-CD45RA (Clone HI100, BioLegend, Cat. No. 304120); anti-CD45RO (Clone UCHL1, BD Biosciences, Cat. No. 560607); anti-CCR7 (Clone 150503, BD Biosciences, Cat. No. 560816); anti-CD69 (Clone FN50, BioLegend, Cat. No. 310936); anti-CD20 (Clone H299, Beckman Coulter, Cat. No. 6604106); anti-CD16 (Clone 3G8, BioLegend, Cat. No. 302016); anti-CD14 (Clone M5E2, BioLegend, Cat. No. 301839); anti-CD11b (Clone M1/70, BD Biosciences, Cat. No. 569708); anti-CD11c (Clone 3.9, BD Biosciences, Cat. No. 565806); anti-HLA-DR (Clone L243, BioLegend, Cat. No. 307640). Following washing, flow cytometry and cell sorting were performed on a BD FACSymphony S6 cell sorter equipped with a 100µm nozzle. Data were acquired using FACSDiva software and subsequently analyzed using FlowJo software (version 10.10.2). Live CD8+ T cells from brain, blood-derived PBMCs, and dura mater were sorted and processed for single-cell RNA sequencing.

### Marmoset CD8^+^ T cell intracellular staining

To measure cytokine expression in CD8^+^ T cells, single-cell suspensions from brain, spleen, blood-derived PBMCs, and dura mater were stimulated with 50 ng/ml PMA and 500 ng/ml ionomycin in the presence of Brefeldin A (3 µg/ml, Invitrogen, Cat. No. 00-4506-51) for 4 hours at 37 °C in a CO incubator. Cells were cultured in complete RPMI 1640 supplemented with 10% FBS, 100 U/ml penicillin, 100 µg/ml streptomycin, 0.25 µg/ml amphotericin B (Invitrogen, Cat. #15240062), and 55 µM β-mercaptoethanol (Thermo Fisher Scientific, Cat. #21985023). Following viability and surface staining, cells were fixed and permeabilized using BD Cytofix/Cytoperm Fixation/Permeabilization Solution (BD Biosciences, Cat. #554714) for 20 minutes at 4 °C. Cells were washed twice with the permeabilization buffer provided and stained for IFN-γ for 30 minutes at 4 °C (anti-human IFN-γ antibody, clone B27, BioLegend Cat. #506516). To measure granzyme B expression without prior stimulation, surface-stained cells were fixed and permeabilized using BD Cytofix/Cytoperm Fixation/Permeabilization Solution (BD Biosciences, Cat. #554714) for 30 minutes at 4 °C. Cells were washed twice with the permeabilization buffer provided and then stained with anti-human granzyme B antibody (clone GB11, BD Biosciences Cat. #563389) for 1 hour at 4 °C.

### Dual-guide CRISPRi lentiviral library

A custom pooled dual-guide CRISPRi lentiviral library targeting 107 surface receptor genes and including NTC sgRNAs was obtained from Sigma-Aldrich (MilliporeSigma; product no. 3009536428). Each gene-targeting construct encoded two sgRNAs targeting the same gene, expressed from separate U6 promoters, and a TagBFP reporter to identify transduced cells. NTC constructs were included to assess effects unrelated to targeted gene repression. The vector incorporated guide-capture sequences compatible with 10x Genomics single-cell analysis. The viral preparation had a supplier-reported physical titer of 4.8 × 10^9^ viral particles/ml, measured by p24 antigen ELISA.

### Mouse primary CD8^+^ T-cell culture, lentiviral transduction, and brain lysate treatment for CRISPR perturb sequencing

On Day 0, primary CD8^+^ T cells were isolated from adult male mouse spleens and mesenteric lymph nodes using the EasySep Mouse CD8^+^ T Cell Isolation Kit (STEMCELL Technologies, Cat. #19853). CD8^+^ T cells were plated at 1×10^5^ cells per well in 200 µl of complete RPMI 1640 supplemented with 10% FBS, 100 U/ml penicillin, 100 µg/ml streptomycin, 0.25 µg/ml amphotericin B (Invitrogen, Cat. #15240062), and 55 µM β-mercaptoethanol (Thermo Fisher Scientific, Cat. #21985023). Plates were coated with Ultra-LEAF Purified anti-mouse CD3ε (BioLegend, Cat. #100340) at 3 µg/ml and Ultra-LEAF Purified anti-mouse CD28 (BioLegend, Cat. #102116) at 6 µg/ml. Cells were plated under identical conditions and assigned to three experimental conditions consisting of lentivirus with brain lysate, lentivirus without brain lysate, and an untransduced control without brain lysate. On Day 1, all cells in the virus-treated groups received 20 µl of medium containing 0.5 µl of our pooled lentiviral CRISPRi library, corresponding to 2.4×10 viral particles per well based on the supplier-reported p24 ELISA titer. On Day 2, virus-containing medium was removed and replaced with fresh complete medium, with brain lysate added to the indicated group. Brain lysate was prepared from mice following transcardial perfusion with 30 mL of cold PBS. Brains were homogenized in complete RPMI 1640 supplemented with Protease Inhibitor Cocktail (Sigma-Aldrich, Cat. #11873580001) at 40 µl of medium per mg of brain tissue. Homogenates were centrifuged at 600 × g for 5 minutes at 4 °C, and the supernatants were collected and filtered through a 0.45-µm filter before addition to CD8^+^ T-cell cultures. Each lysate-treated well received 40 µl of brain lysate in a final culture volume of 200 µl, equivalent to 1 mg of brain tissue per well. On Day 3.5, cells were collected and stained with the LIVE/DEAD Fixable Near-IR Dead Cell Stain Kit (Invitrogen, Cat. #L34993). Live BFP^+^ CD8^+^ T cells were isolated from the lentivirus-treated groups by fluorescence-activated cell sorting with cytometric analysis using BD FACSymphony S6 sorter and processed for library preparation for single-cell RNA sequencing (10x Genomics).

### Single cell RNA sequencing library preparation

Single-cell RNA-sequencing libraries were prepared using the 10x Genomics platform according to the manufacturer’s instructions. Briefly, sorted live BFP CD8 T cells or T cells from marmoset tissues were processed to generate single-cell suspensions and loaded onto the 10x Genomics Chromium system for single-cell partitioning and barcoding. Individual cells were encapsulated in gel bead-in-emulsions (GEMs), where cellular mRNA was captured and reverse-transcribed into cDNA with cell-specific barcodes and unique molecular identifiers (UMIs). The resulting cDNA was amplified and used to construct sequencing libraries according to the manufacturer’s protocol. The final libraries were quantified and assessed for quality before sequencing on a compatible next-generation sequencing platform.

### Single cell RNA sequencing data processing and cross-species integration

Mouse scRNA-seq reference datasets were reanalyzed from our previous study.^31^ To enable direct cross-species transcriptomic comparisons, we processed single-cell RNA sequencing datasets derived from peripheral blood, brain parenchyma, and dura/meninges of common marmosets (*Callithrix jacchus*) in R (v.4.5.3) using Seurat (v.5.5.1). To normalize gene expression profiles while modeling cell-specific sequencing depth, we normalized raw counts independently for mouse and marmoset cells using regularized negative binomial regression via Seurat’s SCTransform function in sctransform (v.0.4.3). To identify shared transcriptomic correspondences across species, we computed cross-species transfer anchors in the SCT space using canonical correlation analysis (CCA) via the FindTransferAnchors function. We then assigned lineage annotations (CD8^+^ T cells, T cells, and NKT cells) to marmoset query cells using the TransferData function. Within each lineage cohort, we split the mouse and marmoset datasets by species and re-normalized them independently with SCTransform. To capture shared biological variance across species, we selected the top conserved variable integration features using the SelectIntegrationFeatures function. We harmonized Pearson residuals across datasets using the PrepSCTIntegration function and identified cross-species integration anchors using the FindIntegrationAnchors function. To project mouse and marmoset cells into a unified, batch-corrected expression space while preserving unintegrated normalized values in the RNA and SCT count layers for downstream differential expression testing, we performed cross-species integration using the IntegrateData function. Principal component analysis was computed on the integrated SCT assay using the RunPCA function, followed by shared nearest-neighbor (SNN) graph construction using the FindNeighbors function and two-dimensional visualization using the RunUMAP function. Cell clustering was performed using the FindClusters function across multiple resolutions.

### Pseudotime trajectory inference and continuous dynamic modeling

To reconstruct developmental transitions and tissue-migration dynamics of purified marmoset CD8^+^ T cells, we performed pseudotime trajectory inference using Monocle 2 (v.2.38.0). We identified high-dispersion ordering genes using the estimateDispersions function, constructed trajectory graphs using DDRTree via reduceDimension, and ordered cells along pseudotime rooted at the blood-predominant naive/central memory branch (state 1) using the orderCells function. To identify dynamically regulated genes across pseudotime and trajectory states, we performed likelihood ratio tests using the differentialGeneTest function, grouped significant genes into four co-expression modules, and evaluated Gene Ontology enrichment using the enrichGO function in clusterProfiler (v.4.18.4). To model continuous expression kinetics without binning or rounding artifacts, we fitted negative binomial generalized additive models directly to raw UMI counts with library-size offsets using tradeSeq (v.1.24.0) via the fitGAM function. We evaluated pseudotime dynamic expression and tissue-specific divergence using the associationTest and conditionTest functions, respectively, and extracted smooth predicted expression curves using the predictSmooth function. We also fitted penalized cubic regression splines using mgcv (v.1.9.4) to confirm continuous dynamics across normalized expression layers.

### Gene regulatory network inference and regulon analysis

To identify transcription factor regulatory networks active in marmoset CD8^+^ T cells, we built a custom *Callithrix jacchus* cisTarget database and applied pySCENIC. We mapped human motif annotations to marmoset orthologs via Ensembl BioMart and extracted promoter sequences from the marmoset reference genome (mCalJac1.pat.X) using bedtools getfasta. We scored promoter regions against vertebrate position weight matrices using Cluster-Buster to generate a gene-by-motif ranking database. To reconstruct regulons, we inferred co-expression modules from purified marmoset CD8^+^ T cells using GRNBoost2 via the pyscenic grn command and pruned indirect targets based on motif enrichment using the pyscenic ctx command, which yielded 47 high-confidence regulons. To quantify regulon activity at single-cell resolution, we computed enrichment scores using the pyscenic aucell command, imported the activity matrix into Seurat, and identified differentially active regulons across trajectory states and tissues using two-sided Wilcoxon rank-sum tests via the FindMarkers function.

### Single-cell Perturb-seq alignment, protospacer calling, and quality control

To quantify single-cell transcriptomes and sgRNA captures simultaneously, we processed paired-end 3′ Gene Expression (GEX) and CRISPR Guide Capture libraries using 10x Genomics Cell Ranger Multi (v.10.1.0) and aligned reads to the mouse reference transcriptome (refdata-gex-mm10-2020-A, GRCm38/mm10). To filter out low-complexity droplets and damaged cells, we performed quality control in Scanpy (v.1.10) and Seurat (v.5.5.1).

### Differential clonal fitness and on-target knockdown evaluation

To determine whether specific gene disruptions confer selective survival or proliferative advantages under BL conditioning, we quantified differential clonal fitness between BL and VO. We then calculated the relative enrichment of each target gene normalized to internal NTC representation. To evaluate target mRNA reduction, we compared target gene expression between perturbed cells and condition-matched NTC controls to compute percentage knockdown efficiency and log2 fold-change on-target values.

### Perturbation signature modeling and functional knockout classification

To distinguish authentic functional knockout (KO) cells from non-perturbed (NP) escapees that failed to undergo gene ablation, we applied the Mixscape framework in Seurat (v.5.5.1) and Scanpy. To isolate guide-specific transcriptional responses from baseline cell-cycle and microenvironmental heterogeneity, we computed local perturbation signatures (PRTB assay) using the CalcPerturbSig function. To visualize low-dimensional phenotypic divergence among knockouts, we performed supervised Linear Discriminant Analysis on validated KO cells using the MixscapeLDA function and projected discriminant vectors onto a post-LDA UMAP manifold. Within this space, we calculated the Euclidean divergence distance of each knockout centroid from the unperturbed NTC centroid and quantified cross-condition state-transition vectors induced by brain lysate exposure.

### Transcriptomic profiling of unperturbed baseline states and microenvironmental shifts

To establish the baseline immunological impact of brain lysate conditioning independent of genetic perturbation, we analyzed unperturbed cohorts (NTC). We performed joint single-cell integration using single-cell Variational Inference (scVI) and Seurat v5. We performed graph clustering with the Leiden algorithm (resolution = 0.45), which resolved resting effector memory, activated cytotoxic, and quiescent/naive-like baseline states.

To identify transcriptome-wide alterations induced by brain lysate, we performed two-sided Wilcoxon rank-sum tests and conducted gene set enrichment analysis (GSEA) against MSigDB Hallmark and Gene Ontology Biological Process (GO:BP) databases using clusterProfiler (v.4.18.4) and gseapy. We calculated single-cell functional module scores using the AddModuleScore function in Seurat and scanpy.tl.score_genes across cytotoxic, inflammatory cytokine, terminal effector TF, naive/memory, and activation/checkpoint programs.

### Systematic effector regulation mapping and two-dimensional perturbation planes

To determine how each genetic perturbation alters CD8^+^ T-cell effector output under brain lysate exposure, we analyzed a 10-gene readout panel (*Gzmb*, *Gzma*, *Prf1*, *Ifng*, *Tnf*, *Xcl1*, *Ccl4*, *Ccl5*, *Tcf7*, and *Pdcd1*) across all 107 active targets in BL relative to the BL NTC baseline. We computed relative Z-scores, deltalog(1+x) shifts, and log2FC values paired with two-sided Mann–Whitney U tests. We generated hierarchical and *Gzmb*-gradient-sorted heatmaps to rank perturbations by cytolytic output.

To systematically evaluate how candidate gene disruptions alter effector and receptor expression across microenvironments, we constructed two-dimensional cross-condition expression planes comparing mean log-normalized expression in brain lysate-treated cells (x-axis) against virus-only cells (y-axis). We scaled bubble areas proportionally to total cell recovery across both conditions and colored circles using a diverging colormap centered at zero to encode the cross-condition differential.

## Resource Availability

Lead contact: Further information and requests for resources and reagents should be directed to and will be fulfilled by the lead contact, Yeong Shin Yim.

## Acknowledgements

We thank members of the Yim and Hildebrand laboratories for technical assistance and discussions. We would like to thank Erich D. Jarvis for providing animals, Alejandra Gonzalez for animal support, and Winrich A. Freiwald for use of facilities and equipment. This work was supported by the Suh Kyungbae Foundation (Y.S.Y.), the NIH through award 1R01MH138484-01A1 (Y.S.Y.), and by start-up funds from the University of Houston Division of Research and College of Optometry (D.G.C.H.).

## Author Contribution

Designed the experiments and/or provided advice and technical expertise, K.P, T.S., S.J., D.G.C.H., and Y.S.Y. Immune cell profiling experiments, K.P. and T.S. scRNA-sequencing and CRISPRi screening Library preparation and bioinformatic analysis, S.J. Marmoset tissues collection P.F.S. and D.G.C.H. Manuscript preparation, K.P, T.S., S.J., D.G.C.H., and Y.S.Y.

## REFERENCES

1. Zinkernagel, R. M. & Doherty, P. C. Restriction of in vitro T cell-mediated cytotoxicity in lymphocytic choriomeningitis within a syngeneic or semiallogeneic system. Nature 248, 701–702 (1974).

2. Murali-Krishna, K. et al. Counting antigen-specific CD8 T cells: a reevaluation of bystander activation during viral infection. Immunity 8, 177–187 (1998).

3. Zhang, N. & Bevan, M. J. CD8(+) T cells: foot soldiers of the immune system. Immunity 35, 161– 168 (2011).

4. Kumar, B. V. et al. Human tissue-resident memory T cells are defined by core transcriptional and functional signatures in lymphoid and mucosal sites. Cell Rep. 20, 2921–2934 (2017).

5. Schenkel, J. M. & Pauken, K. E. Localization, tissue biology and T cell state - implications for cancer immunotherapy. Nat. Rev. Immunol. 23, 807–823 (2023).

6. Casey, K. A. et al. Antigen-independent differentiation and maintenance of effector-like resident memory T cells in tissues. J. Immunol. 188, 4866–4875 (2012).

7. Mackay, L. K. et al. The developmental pathway for CD103(+)CD8+ tissue-resident memory T cells of skin. Nat. Immunol. 14, 1294–1301 (2013).

8. Pan, Y. et al. Survival of tissue-resident memory T cells requires exogenous lipid uptake and metabolism. Nature 543, 252–256 (2017).

9. Masopust, D. & Schenkel, J. M. The integration of T cell migration, differentiation and function. Nat. Rev. Immunol. 13, 309–320 (2013).

10. Masopust, D., Vezys, V., Marzo, A. L. & Lefrançois, L. Preferential localization of effector memory cells in nonlymphoid tissue. Science 291, 2413–2417 (2001).

11. Steinert, E. M. et al. Quantifying memory CD8 T cells reveals regionalization of immunosurveillance. Cell 161, 737–749 (2015).

12. Rustenhoven, J. et al. Functional characterization of the dural sinuses as a neuroimmune interface. Cell 184, 1000–1016.e27 (2021).

13. Groh, J. et al. Accumulation of cytotoxic T cells in the aged CNS leads to axon degeneration and contributes to cognitive and motor decline. Nat. Aging 1, 357–367 (2021).

14. Sauer, B. M., Schmalstieg, W. F. & Howe, C. L. Axons are injured by antigen-specific CD8(+) T cells through a MHC class I- and granzyme B-dependent mechanism. Neurobiol. Dis. 59, 194–205 (2013).

15. Smyth, L. C. D. & Kipnis, J. Redefining CNS immune privilege. Nat. Rev. Immunol. 25, 766–775 (2025).

16. Medawar, P. B. Immunity to homologous grafted skin; the fate of skin homografts transplanted to the brain, to subcutaneous tissue, and to the anterior chamber of the eye. Br. J. Exp. Pathol. 29, 58– 69 (1948).

17. Murphy, J. B. & Sturm, E. Conditions determining the transplantability of tissues in the brain. J. Exp. Med. 38, 183–197 (1923).

18. Castellani, G., Croese, T., Peralta Ramos, J. M. & Schwartz, M. Transforming the understanding of brain immunity. Science 380, eabo7649 (2023).

19. Shirai, Y. On the transplantation of the rat sarcoma in adult heterogeneous animals. The Japan Medical World 1, 14–15 (1921).

20. Wakim, L. M., Woodward-Davis, A. & Bevan, M. J. Memory T cells persisting within the brain after local infection show functional adaptations to their tissue of residence. Proc. Natl. Acad. Sci. U. S. A. 107, 17872–17879 (2010).

21. Huseby, E. S. et al. A pathogenic role for myelin-specific CD8(+) T cells in a model for multiple sclerosis. J. Exp. Med. 194, 669–676 (2001).

22. Chen, X. et al. Microglia-mediated T cell infiltration drives neurodegeneration in tauopathy. Nature 615, 668–677 (2023).

23. Ritzel, R. M. et al. Age-associated resident memory CD8 T cells in the central nervous system are primed to potentiate inflammation after ischemic brain injury. J. Immunol. 196, 3318–3330 (2016).

24. Smolders, J. et al. Tissue-resident memory T cells populate the human brain. Nat. Commun. 9, 4593 (2018).

25. Hsiao, C.-C. et al. Acquisition of residency programs by T cells entering the human brain. Cell Rep. 44, 115960 (2025).

26. Pasciuto, E. et al. Microglia require CD4 T cells to complete the fetal-to-adult transition. Cell 182, 625–640.e24 (2020).

27. Yoshida, T. M. et al. The subfornical organ is a nucleus for gut-derived T cells that regulate behaviour. Nature 643, 499–508 (2025).

28. Korin, B. et al. High-dimensional, single-cell characterization of the brain’s immune compartment. Nat. Neurosci. 20, 1300–1309 (2017).

29. Filiano, A. J. et al. Unexpected role of interferon-γ in regulating neuronal connectivity and social behaviour. Nature 535, 425–429 (2016).

30. Derecki, N. C. et al. Regulation of learning and memory by meningeal immunity: a key role for IL-4. J. Exp. Med. 207, 1067–1080 (2010).

31. Park, K. et al. Brain-resident CD8^+^ T cells regulate neuronal activity and behavior via interferon-gamma. bioRxiv 2026.08.24.746768 (2026) doi:10.64898/2026.08.24.746768.

32. Heusel, J. W., Wesselschmidt, R. L., Shresta, S., Russell, J. H. & Ley, T. J. Cytotoxic lymphocytes require granzyme B for the rapid induction of DNA fragmentation and apoptosis in allogeneic target cells. Cell 76, 977–987 (1994).

33. Trapani, J. A. & Smyth, M. J. Functional significance of the perforin/granzyme cell death pathway. Nat. Rev. Immunol. 2, 735–747 (2002).

34. Dulken, B. W. et al. Single-cell analysis reveals T cell infiltration in old neurogenic niches. Nature 571, 205–210 (2019).

35. Brizić, I. et al. Brain-resident memory CD8+ T cells induced by congenital CMV infection prevent brain pathology and virus reactivation. Eur. J. Immunol. 48, 950–964 (2018).

36. Liu, Y., Teige, I., Birnir, B. & Issazadeh-Navikas, S. Neuron-mediated generation of regulatory T cells from encephalitogenic T cells suppresses EAE. Nat. Med. 12, 518–525 (2006).

37. Chen, Y., Menegas, W., Zhang, Q. & Feng, G. Common marmoset: An emerging non-human primate model for translational applications in brain disorders. Curr. Opin. Neurobiol. 92, 102998 (2025).

38. Hebbar, P. et al. A complete genome for the common marmoset. Cell 189, 4891–4908.e12 (2026).

39. Jagessar, S. A. et al. Overview of models, methods, and reagents developed for translational autoimmunity research in the common marmoset (Callithrix jacchus). Exp. Anim. 62, 159–171 (2013).

40. Zarif, H. et al. CD8+ T cells are essential for the effects of enriched environment on hippocampus-dependent behavior, hippocampal neurogenesis and synaptic plasticity. Brain Behav. Immun. 69, 235–254 (2018).

41. Simon, M. M. et al. In vitro- and ex vivo-derived cytolytic leukocytes from granzyme A x B double knockout mice are defective in granule-mediated apoptosis but not lysis of target cells. J. Exp. Med. 186, 1781–1786 (1997).

42. Weinreb, C., Wolock, S., Tusi, B. K., Socolovsky, M. & Klein, A. M. Fundamental limits on dynamic inference from single-cell snapshots. Proc. Natl. Acad. Sci. U. S. A. 115, E2467–E2476 (2018).

43. Beura, L. K. et al. Normalizing the environment recapitulates adult human immune traits in laboratory mice. Nature 532, 512–516 (2016).

